# Haplotype-rich *cis*-regulation underlies transcriptomic diversity across the breeding history of maize (*Zea mays*)

**DOI:** 10.64898/2026.02.19.706772

**Authors:** Marcin W. Grzybowski, James C. Schnable

## Abstract

The architecture of gene-expression variation and its relationship to population differentiation remain incompletely characterized in crops. Here, we integrated leaf RNA-seq and genome-wide variation from 631 maize inbred lines to characterize transcriptomic diversity and fine-map *cis*-regulatory signals. Subpopulation identity explained a mean of 5.3% of expression variation across genes, indicating that most expression diversity segregates within rather than among breeding populations. We identified 17,468 genes with *cis*-eQTLs and resolved putatively causal eQTLs for 16,270 of them. Most fine-mapped eGenes (77.7%) were haplotype-rich (at least two credible sets), and approximately 39% of credible sets contained a single variant. Modeling all credible sets increased median *R*^2^ from 0.16 to 0.27. Individual effects were generally modest, and genes with more signals tended to have higher expression heritability. Fine-mapped variants were enriched in transcribed regions, accessible chromatin, and transcription-factor-binding regions, with the latter enrichment persisting after accessibility matching. Genes with high gene-body sequence constraint had modestly smaller regulatory effects and fewer credible sets. Across five population contrasts, including Stiff Stalk–Non-Stiff Stalk comparisons within two breeding eras, genes with stronger statistical evidence of differential expression were associated with higher *F*_ST_ at fine-mapped variants. These results indicate that single-variant summaries provide an incomplete picture of *cis*-regulatory loci, and offer a mechanism for why transcriptome-wide association studies identify genes missed by genome-wide association studies in the same populations.

## Introduction

Genetic variation influences phenotype through multiple molecular layers, among which gene expression represents a primary and highly dynamic intermediate. Variation in transcript abundance across individuals can be associated with specific DNA variants, commonly referred to as expression quantitative trait loci (eQTLs). Regulatory variation is thought to play a central role in evolution and adaptation, as changes in gene regulation can act in a tissue- or context-specific manner and may be less constrained by pleiotropic effects than mutations affecting protein-coding sequences. Consistent with this view, variation in non-coding regions has been estimated to account for a substantial fraction of phenotypic variation in complex traits (Finucane *et al*., 2015; Rodgers-Melnick *et al*., 2016).

Maize (*Zea mays ssp. mays*) is both a major global crop and a powerful model for studying the evolution of regulatory variation. Domesticated approximately 10,000 years ago in Mexico from its wild progenitor, teosinte (*Zea mays* ssp. *parviglumis*) (Matsuoka *et al*., 2002; Yang *et al*., 2023), maize subsequently spread across a wide range of temperate and tropical environments. Since the mid-twentieth century, maize production has been dominated by single-cross hybrid varieties derived from inbred lines belonging to distinct heterotic groups. In the United States, breeding programs have historically relied on three major heterotic groups: the female Stiff Stalk Synthetic (SS) group and the male Non-Stiff Stalk (NSS) and Iodent (IDT) groups (Mikel and Dudley, 2006). Recurrent selection within these groups, combined with limited gene flow between them, has generated pronounced genetic differentiation over successive breeding eras (Li *et al*., 2022).

The genetic and phenotypic consequences of modern maize breeding have been extensively studied at the morphological and genomic levels (Gage *et al*., 2018; Jiao *et al*., 2012; Li *et al*., 2022; Unterseer *et al*., 2016; van Heerwaarden *et al*., 2012; Wang *et al*., 2020a). In contrast, considerably less attention has been paid to how breeding and population structure have shaped the maize transcriptome. Recent work has begun to address this gap, including a study of Chinese maize inbred lines that reported widespread expression differences consistent with selection acting on regulatory variation (Li *et al*., 2023). However, the extent to which transcriptomic diversity mirrors genomic diversity across heterotic groups remains poorly understood.

Numerous studies have mapped eQTLs in maize using recombinant populations or diversity panels (Kremling *et al*., 2018; Liu *et al*., 2017, 2020; Sun *et al*., 2023a; Wang *et al*., 2018). Recombinant populations provide substantial power for eQTL detection, but their extended linkage disequilibrium limits mapping resolution. Diversity panels offer finer resolution because of more extensive historical recombination, yet eQTLs are still commonly represented by a single lead variant. Such representations can obscure allelic heterogeneity, in which multiple variants or regulatory haplotypes independently contribute to variation in expression of the same gene. Statistical fine-mapping approaches can distinguish multiple association signals and define credible sets of variants likely to contain the underlying regulatory variants (Li and Zhou, 2025). Studies in humans have revealed widespread allelic heterogeneity of *cis*-regulatory effects (Taylor *et al*., 2024), but the prevalence and structure of such complex regulatory architectures remain much less characterized in plants.

Here, we use RNA-seq and dense genome-wide genetic variation from 631 diverse maize inbred lines to investigate how population and breeding history are reflected in transcriptomic diversity and its genetic architecture. We first quantify differences in genomic and transcriptomic diversity among major maize subpopulations. We then map and fine-map *cis*-eQTLs to determine the extent of allelic heterogeneity underlying gene-expression variation and characterize the genomic distribution and effect sizes of fine-mapped regulatory variants. Finally, we examine whether expression-associated variants show altered genetic differentiation between different maize subpopulation and breading eras. Together, these analyses link population-level transcriptomic variation to the fine-scale, haplotype-rich architecture of *cis*-regulatory variation in modern maize.

## Results

### Transcriptomic diversity across maize subpopulations

We analyzed RNA-seq data from 631 maize inbred lines sampled in the field within a two-hour interval on the same day. Samples were collected from the preantepenultimate leaf when approximately 50% of the inbreds had reached flowering (Torres-Rodríguez *et al*., 2024). The lines belonged to the Wisconsin Diversity Panel (WiDiv) (Mazaheri *et al*., 2019) and had previously been whole-genome resequenced (Grzybowski *et al*., 2023). Based on previously defined population assignments (Mazaheri *et al*., 2019), the panel comprised six genetic groups: Broad-origin public (n = 129), Iodent (n = 53), Mixed (n = 148), Non-Stiff Stalk (NSS, n = 114), Stiff Stalk (SS, n = 155), and Tropical (n = 32; Table S1). The Mixed group comprised lines with an Admixture-inferred membership probability *<* 0.5 for each defined subpopulation, whereas the Broad-origin public group consisted of genetically distinct public inbred lines not assigned to the major heterotic groups. Comparison with a broader panel of 1,515 maize accessions showed that the WiDiv lines spanned much of the genetic diversity represented in modern maize (Fig. S4).

PCA of genome-wide genetic variation showed pronounced population structure, particularly among the three major U.S. heterotic groups: SS, NSS, and Iodent (Mikel and Dudley, 2006). PC1 primarily separated SS from NSS and Iodent, whereas PC2 distinguished Iodent from NSS (Fig. 1A, Table S1), consistent with previous analyses of North American maize germplasm (van Heerwaarden *et al*., 2012; Wang *et al*., 2020a).

**Figure 1.**
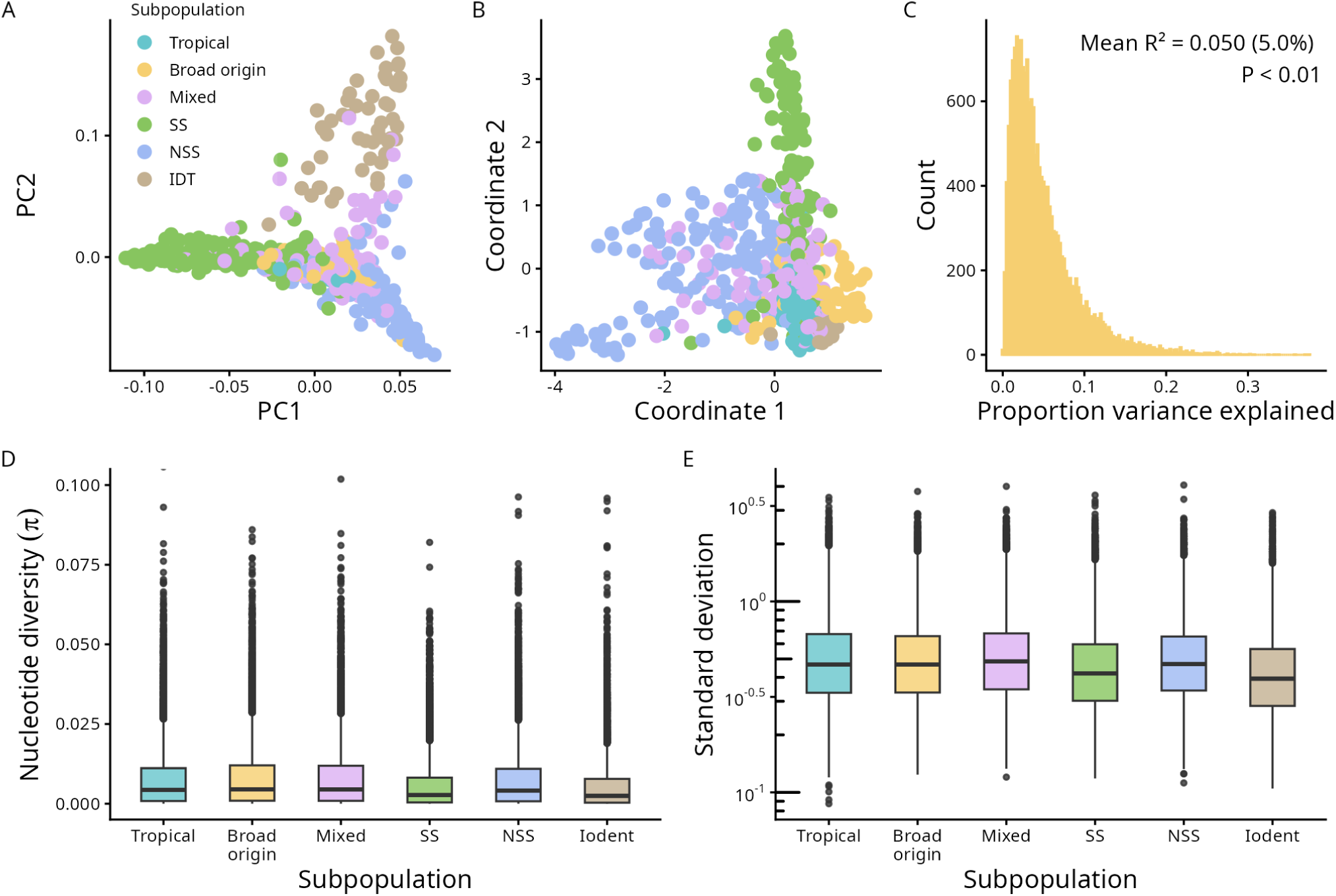
Genomic and transcriptomic diversity across maize subpopulations. (**A**) Principal component analysis (PCA) based on genome-wide genetic variants, showing clear separation of major maize heterotic groups. (**B**) Canonical correlation analysis (CCA) projecting gene expression variation onto the genetic PCA space, revealing detectable but weaker population structure in transcriptomic data. (**C**) Proportion of variance in flowering-time-adjusted gene expression explained by subpopulation identity. (**D**) Nucleotide diversity (*π*) across maize subpopulations (permutation ANOVA, *P <* 1 *×* 10*^−^*^16^). (**E**) Transcriptomic diversity, measured as the standard deviation of flowering-time-adjusted, variance-stabilized expression (ANOVA based on mix-model, *P <* 1 *×* 10*^−^*^16^).

To determine whether this genetic population structure was also reflected in gene-expression variation, we used two complementary approaches. First, canonical correlation analysis (CCA) was used to assess concordance between genetic and transcriptomic variation (Brown *et al*., 2018). CCA revealed detectable but weaker differentiation among subpopulations at the transcriptomic level (Fig. 1B, Table S1). The first canonical coordinate primarily distinguished NSS lines from the remainder of the panel, whereas the second separated SS lines from other groups. Second, we quantified the proportion of gene-expression variation associated with subpopulation identity using gene-wise linear models. Subpopulation explained a mean of 5.3% of expression variation across genes (Fig. 1C), significantly more than expected under a null distribution generated by permutation of subpopulation labels (one-tailed permutation test, *P <* 0.01). Together, these analyses showed that genetic population structure is reflected in gene-expression variation, although transcriptomic differentiation is substantially weaker than genomic differentiation.

We next asked whether differences in genomic diversity among subpopulations were accompanied by corresponding differences in transcriptomic diversity. Nucleotide diversity (*π*) differed substantially among genetic groups, with nearly twofold lower diversity in SS (*π* = 0.00270) and Iodent (*π* = 0.00245) than in Tropical lines (*π* = 0.00431; Fig. 1D; permuatation ANOVA, *P <* 1 *×* 10*^−^*^16^). The Tropical lines represented in WiDiv nevertheless contained substantially less nucleotide diversity than the broader set of Tropical accessions in the larger maize diversity panel (median *π* = 0.0105; Grzybowski *et al*., 2023).

Transcriptomic diversity followed the same general ordering among subpopulations, but the magnitude of the differences was considerably smaller. Relative to Tropical lines, transcriptomic diversity was reduced by *∼*10% in SS and *∼*20% in Iodent (Fig. 1E; ANOVA based on mix-model, *P <* 1 *×* 10*^−^*^16^).

### High-resolution fine-mapping of *cis*-regulatory variants

The correspondence between genetic and transcriptomic population structure suggests that expression divergence may arise, at least in part, from differences in allele frequencies at regulatory variants. To characterize this regulatory variation, we mapped *cis*-acting expression quantitative trait nucleotides (eQTNs) by integrating gene expression with previously published genotype data for the same maize inbred lines (Grzybowski *et al*., 2023). Variants located within 2 Mb of each gene’s transcription start site (TSS) were tested for association with expression using a mixed linear model. We defined eGenes as genes with at least one significant eQTN. Among the 18,197 expressed genes that passed our filtering criteria, 17,468 (96%) were classified as eGenes. In total, we identified 15,101,889 unique eQTNs and 69,297,277 significant eGene-eQTN pairs at a 1% false discovery rate (FDR), corresponding to a mean of 3,967 and a median of 3,156 associated variants per eGene (Fig. S5A). Significant eQTNs were concentrated near the TSS of their associated genes (Fig. S5B). Their minor allele frequency (MAF) distribution was consistent with previous eQTL studies in humans and maize (Mostafavi *et al*., 2023; Sun *et al*., 2023a), with 12.6% of significant eQTNs having MAF *<* 0.05 (Fig. S5C).

Because eQTL discovery is strongly dependent on statistical power, we used the comparatively large sample size of the WiDiv panel to examine how eGene discovery scaled with population size. We repeated the *cis*-eQTL analysis in progressively larger stratified subsets while preserving the relative representation of the major genetic subpopulations. The number of genes with at least one significant *cis*-eQTL increased rapidly at smaller sample sizes and gradually approached an asymptote in the largest subsets. At a sample size of 500 lines, approximately 95% of the estimated asymptotic number of eGenes had been detected (Fig. S6). These results indicate that the high proportion of eGenes identified in the complete panel was enabled by its substantial statistical power, while the continued increase in discovery at larger sample sizes suggests that additional regulatory associations would likely be detected in still larger populations.

The large number of significant eQTNs associated with individual genes highlights an important limitation of association mapping: many significant variants are correlated through linkage disequilibrium (LD) and therefore cannot be interpreted as independent regulatory effects. Although LD decays rapidly in the WiDiv population (LD *<* 0.2 within 2 kb; Grzybowski *et al*., 2023), local LD can still generate large groups of associated variants around individual regulatory signals. We therefore fine-mapped the *cis*-eQTL associations using SuSiE (Wang *et al*., 2020b) to distinguish independent regulatory signals and narrow the set of variants likely to contain their causal variants. SuSiE models the association pattern as a sum of single effects, with each inferred signal represented by a credible set of variants containing the putative causal variant with high posterior probability. At least one credible set was identified for 93% of eGenes (*n* = 16,270; Table S2). Fine-mapping revealed extensive allelic heterogeneity, with 77.7% of successfully fine-mapped eGenes containing two or more independent credible sets (mean = 2.77; median = 2; Fig. 2A). Thus, most eGenes have haplotype-rich regulatory architectures, segregating for multiple regulatory haplotypes rather than two alternative alleles.

**Figure 2.**
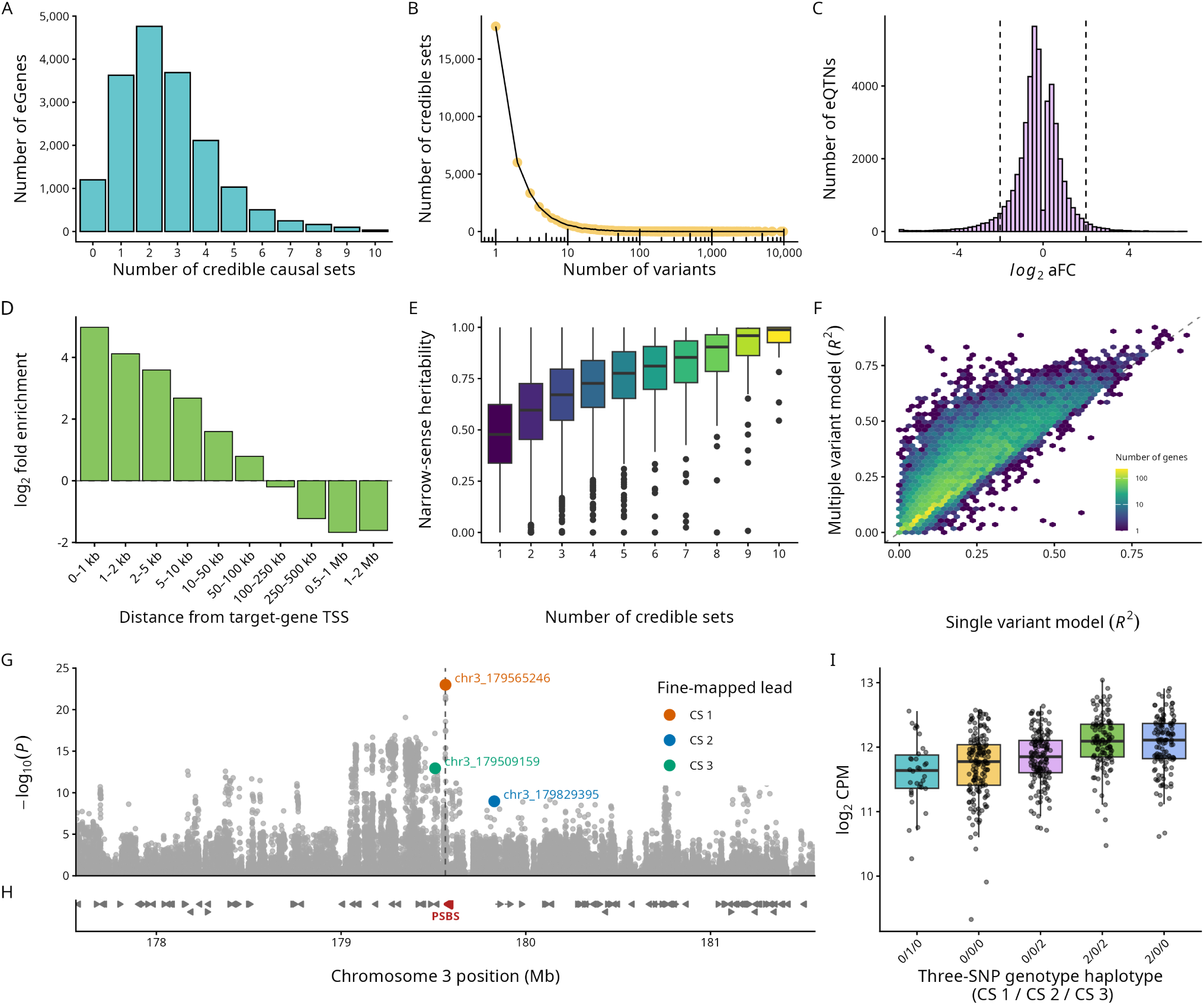
Fine-mapping architecture and a multi-variant eQTL example. (**A**) Number of credible sets per eGene. (**B**) Relationship between the number of variants and the number of credible sets. (**C**) Distribution of allelic fold changes (aFCs) for eQTNs; dashed lines indicate log_2_ aFC values of *−*2 and 2. (**D**) Enrichment of fine-mapped variants by distance from the target-gene transcription start site. (**E**) Narrow-sense heritability by number of credible sets. (**F**) Agreement between single-variant and multiple-variant eQTL-model *R*^2^ values. (**G–H**) Local association signal and gene track for *PSBS* (Zm00001eb146510), with credible-set lead variants highlighted. (**I**) Expression across the most frequent three-SNP haplotypes.

A large fraction of credible sets contained only a single variant (*∼*39%, *n* = 17,839), indicating high mapping resolution (median = 2 variants per credible set; Fig. 2B). We selected the variant with the highest posterior inclusion probability from each credible set and estimated its effect using allelic fold change (Ehsan *et al*., 2024), jointly accounting for other fine-mapped lead eQTNs associated with the same gene. Regulatory effects were generally modest, with a median *|* log_2_(aFC)*|* of 0.59 (Fig. 2C; Table S3). Fine-mapped variants were also most strongly enriched close to the TSS of their target genes, with enrichment declining with increasing TSS distance (Fig. 2D), consistent with a predominantly local architecture of *cis*-regulatory variation.

We next examined whether the complexity of *cis*-regulatory architecture was associated with the genetic contribution to gene-expression variation. Narrow-sense expression heritability was estimated from a genomic relationship matrix. Because expression PCs and other covariates included in the eQTL model capture substantial non-genetic and structured variation, heritability was estimated from both unadjusted and covariate-adjusted expression values (Fig. S7). Covariate adjustment increased median narrow-sense heritability from 0.54 to 0.61 (paired Wilcoxon signed-rank test, *P <* 2.2 *×* 10*^−^*^16^), and covariate-adjusted estimates were used in subsequent analyses.

Expression heritability varied considerably among genes and was related to the complexity of their fine-mapped regulatory architecture. Genes containing a larger number of independent credible sets tended to have higher narrow-sense heritability (Fig. 2E). We additionally examined whether this relationship could be explained by basic expression characteristics, including mean expression and expression variance. Although these properties varied among genes, the association between regulatory complexity and expression heritability remained evident (Fig. S8). Together, these results indicate that genes with a larger genetically determined component of expression variation tend to harbor a more complex *cis*-regulatory architecture.

Given the relationship between regulatory complexity and expression heritability, we next asked whether explicitly modeling multiple *cis*-regulatory signals improved prediction of gene expression. We compared a single-variant model containing the most significant marginal eQTN with a multi-variant model containing one lead variant from each SuSiE credible set. Model coefficients were estimated in a training subset and predictive performance was evaluated in held-out lines. Test-set performance was strongly correlated between the two approaches (Pearson’s *r* = 0.78, *P <* 2.2 *×* 10*^−^*^16^), but the multi-variant model explained substantially more expression variation, increasing median test-set *R*^2^ from 0.16 to 0.27 (paired Wilcoxon signed-rank test, *P<* 2.2 *×* 10*^−^*^16^, Fig. 2F). Improvements were most apparent for genes containing multiple distinct fine-mapped *cis*-regulatory signals (Fig. S9).

To illustrate this multi-signal architecture at an individual locus, we examined *PHOTOSYSTEM II SUBUNIT S* (*PSBS*; Zm00001eb146510), for which we previously identified a *cis*-eQTL associated with variation in gene expression (Sahay *et al*., 2023). The local association profile comprised a broad cluster of significant variants surrounding *PSBS*, which fine-mapping resolved into three distinct regulatory signals represented by separate lead eQTNs (Fig. 2G, H). These variants occurred in several common haplotypes associated with markedly different levels of *PSBS* expression, making *PSBS* a haplotype-rich locus (Fig. 2I). Consistent with the contribution of multiple regulatory signals at this locus, the held-out test-set *R*^2^ increased from 0.15 for the single-variant model to 0.24 when all three fine-mapped lead variants were modeled jointly. Thus, an apparently complex marginal association pattern could be decomposed into a small number of distinct *cis*-regulatory signals whose combined effects contribute substantially to expression variation within the population.

### Fine-mapped *cis*-regulatory variants preferentially localize to functional genomic regions

We next investigated the genomic context of fine-mapped *cis*-regulatory variants. To account for differences in allele frequency and genomic position, enrichment was evaluated relative to matched control variants selected from the same *cis* regions and matched by minor allele frequency, distance from the target-gene TSS, and variant type. Fine-mapped eQTNs were strongly enriched within transcribed regions, with the greatest enrichment observed in 5*^′^* UTRs (1.63-fold), followed by 3*^′^* UTRs (1.55-fold), coding sequences (1.52-fold), and introns (1.20-fold; Fig. 3A). In contrast, promoter regions showed little enrichment, whereas intergenic variants were depleted relative to the matched genomic background. Although intergenic variants represented the largest individual class of fine-mapped eQTNs, their relative depletion indicates that fine-mapped regulatory signals occur disproportionately within gene-associated functional sequence.

**Figure 3.**
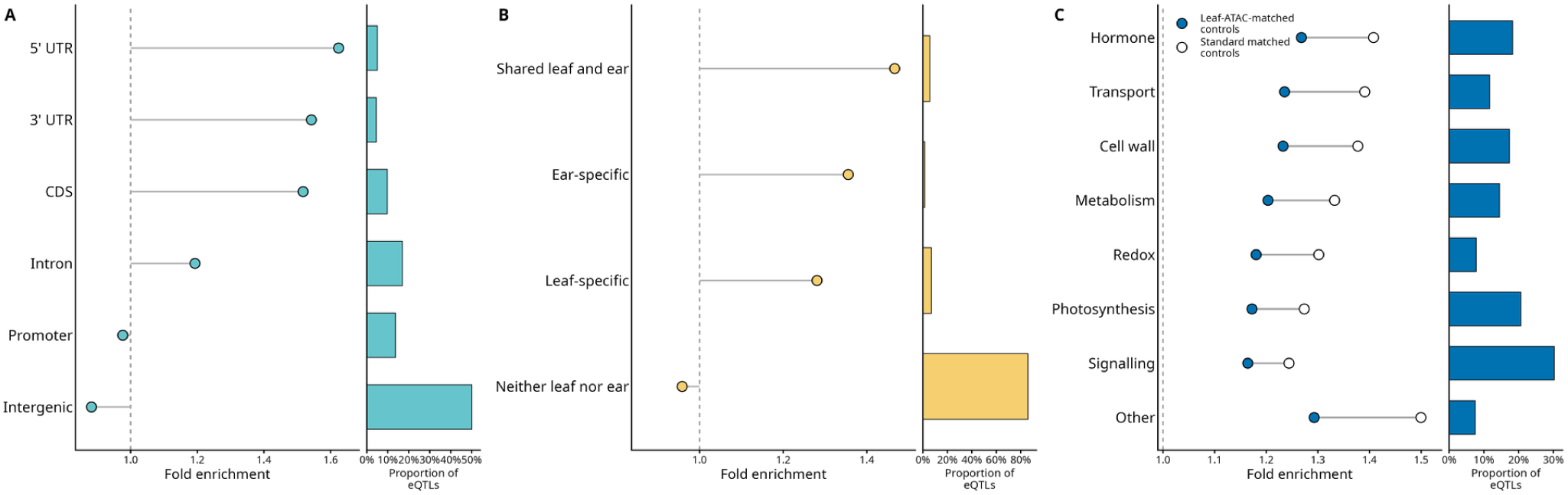
Fine-mapped eQTN enrichment across genomic, tissue-specific, and functional annotations. (**A**) Fold enrichment of fine-mapped eQTNs across genomic features, with the proportion of eQTNs assigned to each feature shown at right. (**B**) Fold enrichment and eQTN proportion by leaf and ear chromatin-accessibility categories. (**C**) Functional-category enrichment of fine-mapped eQTNs using leaf-ATAC-matched and standard matched controls; bars show the proportion of eQTNs in each category.

Fine-mapped variants were also preferentially associated with accessible chromatin (Fig. 3B) (Ricci *et al*., 2019). The strongest enrichment was observed in regions accessible in both leaf and ear tissues (*∼*1.46-fold), followed by ear-specific (*∼*1.35-fold) and leaf-specific (*∼*1.28-fold) accessible regions. Variants outside accessible regions in either tissue showed little or no enrichment. Thus, fine-mapped regulatory variants were preferentially located in open chromatin, with the strongest enrichment occurring in regulatory regions accessible across tissues rather than exclusively in the sampled leaf tissue.

We further asked whether fine-mapped variants preferentially overlapped transcription-factor binding sites and whether this enrichment could be explained simply by their increased occurrence in accessible chromatin (Tu *et al*., 2020). ChIP-seq peaks were grouped according to the functional classes of the corresponding transcription factors, including hormone, transport, cell-wall, metabolism, redox, photosynthesis, and signaling regulators (Fig. 3C). Using controls matched for MAF, TSS distance, and variant type, fine-mapped eQTNs were enriched across all TF functional classes (Fig. S10). Enrichment was reduced when controls were additionally matched for leaf ATAC-seq accessibility, as expected from the strong association between TF binding and open chromatin, but remained above unity across all classes. This persistence after conditioning on chromatin accessibility indicates that the localization of fine-mapped eQTNs within TF-binding regions cannot be explained solely by their preferential occurrence in accessible chromatin and is consistent with direct perturbation of transcriptional regulatory elements.

### Evolutionary constraint limits the magnitude and complexity of *cis*-regulatory variation

If large changes in gene expression are more likely to be deleterious, purifying selection is expected to restrict large-effect regulatory alleles to lower frequencies. Consistent with this expectation, rare regulatory variants have previously been associated with disproportionately large effects on gene expression and expression dysregulation in maize (Kremling *et al*., 2018). In our data, the magnitude of fine-mapped regulatory effects similarly decreased progressively with increasing minor allele frequency (MAF; Fig. 4A). Median *|* log_2_(aFC)*|* decreased from 0.948 among variants with MAF between 0.01 and 0.05 to 0.414 among variants with MAF between 0.40 and 0.50 (Spearman’s *ρ* = *−*0.412, *P <* 2.2 *×* 10*^−^*^16^).

**Figure 4.**
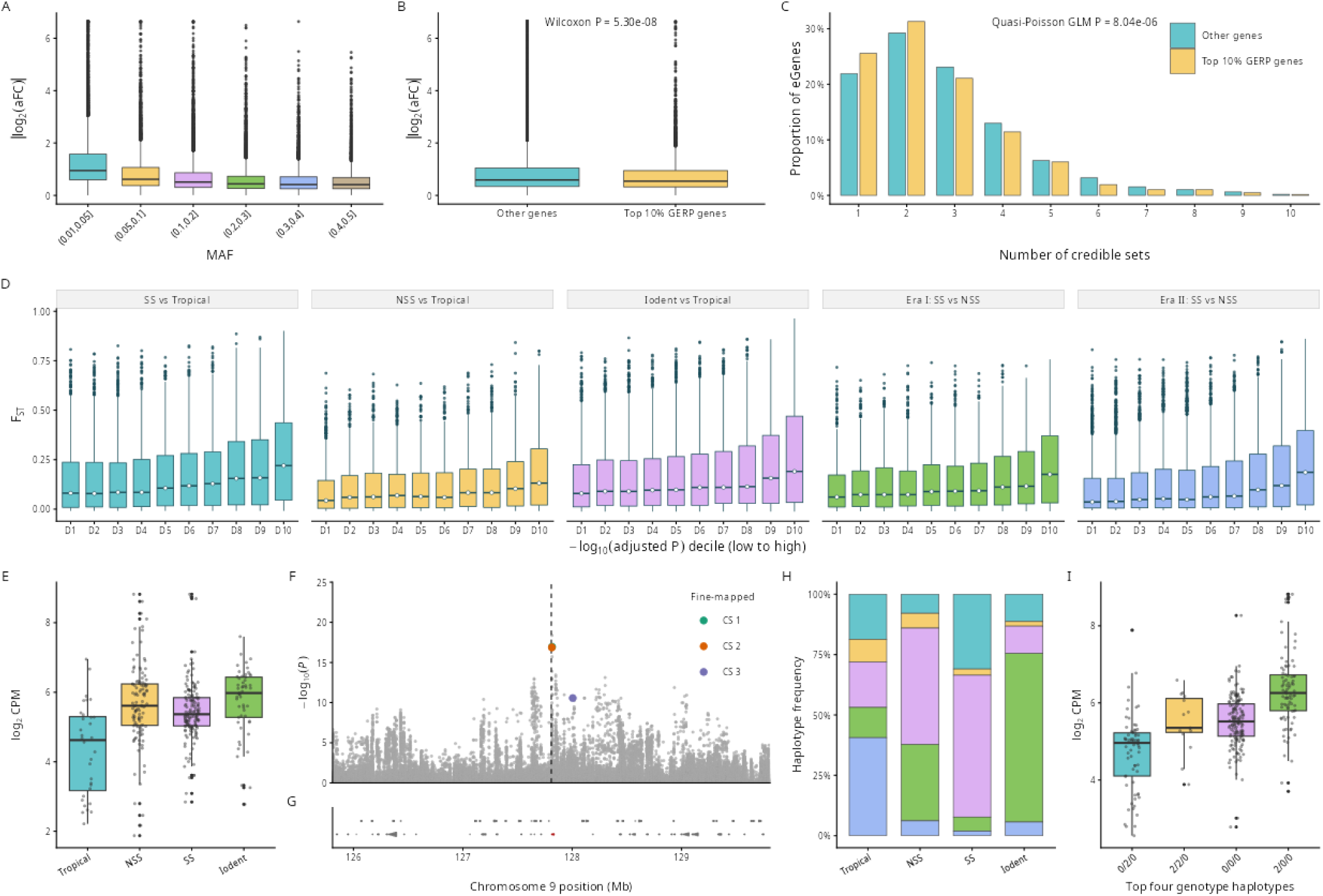
Fine-mapped eQTN effect sizes, population differentiation, and a Zm00001eb392880 example. (**A**) Absolute aFC by minor-allele-frequency bin. (**B**) Absolute aFC for genes in the top decile of positive GERP scores and all other genes. (**C**) Distribution of credible-set counts for the same gene groups. (**D**) *F*_ST_ across adjusted-expression-significance deciles for five population comparisons. (**E**) Zm00001eb392880 expression across Tropical, NSS, SS, and Iodent lines. (**F–G**) Local eQTL association signal and gene track, with fine-mapped credible-set variants highlighted. (**H**) Haplotype frequencies of fine-mapped variants across subpopulations. (**I**) Expression across the four most frequent fine-mapped-variant haplotypes.

We next asked whether this relationship was also reflected in an independent measure of long-term evolutionary constraint. Evolutionary constraint was quantified using Genomic Evolutionary Rate Profiling (GERP) scores (Davydov *et al*., 2010; Hufford *et al*., 2021). Genes in the upper 10% of the distribution of mean positive GERP scores were classified as highly constrained. Fine-mapped eQTNs associated with highly constrained genes had modestly but significantly smaller effects than those associated with the remaining genes (median *|* log_2_(aFC)*|* = 0.544 and 0.590, respectively; Wilcoxon rank-sum test, *W* = 87,538,893, *P* = 5.30 *×* 10*^−^*^8^; Fig. 4B).

Evolutionary constraint was also associated with the complexity of *cis*-regulatory architecture. Highly constrained genes showed a greater proportion of one- and two-signal architectures and a lower frequency of genes containing larger numbers of credible sets (Fig. 4C). The mean number of credible sets was 2.60 for highly constrained genes and 2.78 for the remaining genes (quasi-Poisson generalized linear model, *P* = 8.04 *×* 10*^−^*^6^). Thus, evolutionary constraint was associated with both reduced regulatory effect sizes and a modest reduction in the number of independently fine-mapped *cis*-regulatory signals. This pattern is consistent with purifying selection limiting standing regulatory variation at functionally constrained genes, paralleling patterns reported in humans (Taylor *et al*., 2024).

### Population differentiation at fine-mapped eQTNs accompanies gene-expression divergence

Modern maize population structure has been strongly shaped by breeding history and the development of distinct heterotic groups (Li *et al*., 2022; van Heerwaarden *et al*., 2012). We therefore asked whether population-dependent differences in gene expression were accompanied by allele-frequency differentiation at fine-mapped *cis*-regulatory variants. Differential-expression analyses were restricted to the same set of expressed genes used for eQTL mapping. Five population contrasts were examined: SS versus Tropical, NSS versus Tropical, Iodent versus Tropical, and SS versus NSS within era I and era II germplasm. The number of differentially expressed genes ranged from 4,121 to 10,642 among these contrasts.

Within each contrast, significant differentially expressed genes were ranked according to *−* log_10_-transformed Benjamini–Hochberg-adjusted *p*-values and divided into ten equal-sized groups. Thus, D1 contained genes with the weakest statistical support among significant differentially expressed genes, whereas D10 contained those with the strongest support. Across all five contrasts, *F*_ST_ at fine-mapped eQTNs generally increased across these differential-expression significance deciles (Fig. 4D). Median *F*_ST_ ranged from 0.036–0.081 in D1 and from 0.131–0.220 in D10 across the five comparisons.

The relationship was observed both between Tropical and temperate heterotic groups and between SS and NSS within the era I and era II breeding groups. Thus, genes showing the strongest statistical evidence of population-dependent expression were also more likely to be associated with fine-mapped regulatory variants exhibiting pronounced allele-frequency differentiation. These results are consistent with population-specific shifts in the frequencies of standing *cis*-regulatory alleles contributing to transcriptomic divergence during the formation and subsequent improvement of maize breeding populations (Li *et al*., 2022; van Heerwaarden *et al*., 2012). Although demographic history and linked selection may also contribute to variant differentiation, the consistency of the relationship across multiple contrasts supports a contribution of differentiated regulatory variants to population-level expression differences.

To illustrate this relationship in detail at an individual locus, we examined *SUCROSE SYNTHASE1* (*SUS1*; Zm00001eb392880), which encodes a sucrose-cleaving enzyme involved in carbon partitioning and starch biosynthesis (Chourey *et al*., 1998; Deng *et al*., 2020). *SUS1* expression differed among genetic groups (limma-voom, empirical Bayes-moderated *F*-test for the overall population effect, *P <* 0.01; Fig. 4E). Median expression was 4.62 log_2_-CPM in Tropical lines and 5.98 log_2_-CPM in Iodent. Fine-mapping of the broad local association signal resolved three distinct regulatory signals represented by separate lead eQTNs, identifying *SUS1* as another haplotype-rich locus (Fig. 4F,G). The high-expression three-SNP haplotype (2/0/0) was most frequent in Iodent germplasm (69.8%) and least frequent in Tropical germplasm (12.5%; Fig. 4H). The four most frequent haplotypes were associated with distinct levels of *SUS1*expression (one-way ANOVA, *P <* 2.2 *×* 10*^−^*^16^; Fig. 4I). Together, this locus illustrates how population-specific frequency shifts at multiple standing *cis*-regulatory variants can jointly contribute to expression divergence among maize genetic groups.

## Discussion

In this study, we integrated population-level expression analyses with *cis*-eQTL mapping and fine-mapping to characterize regulatory variation across major maize breeding populations, quantifying two complementary dimensions of transcriptomic diversity: differences in expression profiles among populations and variation among lines within each population. We were able to include substantially more distinct genotypes (631 inbred lines) than several previous maize eQTL panels, which comprised 137–340 accessions (Kremling *et al*., 2018; Li *et al*., 2023; Liu *et al*., 2020; Sun *et al*., 2023b). Approximately 95% of the fitted asymptotic eGene count was reached at 500 lines (Fig. S6), although studies targeting additional signals at already identified eGenes may benefit from further sampling. We found that population-average expression differences captured only part of the transcriptomic variation present in maize breeding germplasm (subpopulation identity explained a mean of 5.3% of expression variation across genes). Characterizing the underlying regulatory architecture therefore requires examining not only variants differentiated among populations, but also the multiple segregating alleles and haplotypes associated with expression variation within them.

In our study of maize, haplotype-rich *cis*-eQTLs frequently consisted of at least two credible sets (at least three haplotypes) corresponding to distinct variants with individually modest effects (median *|* log_2_(aFC)*|* of 0.59) on the expression of the target gene. Incorporating multiple fine-mapped *cis*-eQTNs markedly increased the explained variance relative to single variant approaches. Direct comparisons of *cis*-regulatory architecture across species remain challenging, but, in humans, recent large-scale studies have reported fewer independent credible sets per gene on average than observed here (40% of fine-mapped eGenes with more than one credible set, versus 77.7% here), although widespread multi-signal *cis*-regulation is still evident (Taylor *et al*., 2024). In plants, most eQTL studies rely on single-variant summaries, precluding systematic assessment of allelic heterogeneity at regulatory loci (Li *et al*., 2023; Liu *et al*., 2020; Sun *et al*., 2023a). For these regulatory haplotype-rich (*≥* 3 functionally distinct haplotypes) loci, summarizing a *cis*-eQTL using a single lead variant from the association peak will provide incomplete and potentially misleading information when seeking to link DNA sequence or chromatin polymorphisms to their functional impact on the expression of the target gene. Fine-mapped variants were enriched in transcribed regions and in transcription-factor-binding sites even after matching for chromatin accessibility, consistent with direct perturbation of regulatory elements and providing a basis for prioritizing candidate variants.

Genes with more detectable *cis*-regulatory signals tended to have higher narrow-sense heritability. This association is consistent with a larger genetically determined component of expression variation being accompanied by a greater number of resolvable local effects but is also consistent with our statistical analysis having more power to detect additional signals when a smaller proportion of variation in gene expression is explained by non-genetic factors.

Genes under stronger evolutionary constraint tended to harbor fewer *cis*-eQTNs with smaller effect sizes than genes predicted to be under lower evolutionary constraint, indicating weak purifying selection acting against large regulatory perturbations. The inverse association between effect magnitude and minor allele frequency is compatible with this interpretation, although differential power to detect small effects at rare variants and ascertainment of significant associations may also contribute to that relationship. Comparable trends have been observed in human populations, where regulatory variants at constrained genes typically show attenuated effects on expression (Glassberg *et al*., 2019; Taylor *et al*., 2024). Notably, this constraint operates on the magnitude and combinatorial architecture of regulatory effects rather than on the presence of *cis*-regulatory variation per se, allowing standing regulatory variation to persist while limiting its phenotypic consequences.

The pattern we observed of genes with stronger statistical evidence of differential expression tending to carry fine-mapped variants with higher *F*_ST_ is consistent with differentiated frequencies of segregating *cis*-regulatory alleles contributing to population-level expression differences. Under a multi-signal regulatory model, modest shifts in the frequencies of several modest-effect variants can collectively generate substantial expression differences between heterotic groups, providing a mechanistic link between breeding history, population structure, and transcriptomic divergence. Demographic history and linked selection may also contribute to the observed differentiation. At *SUS1*, fine-mapping resolved three signals, and the haplotype associated with higher expression was substantially more frequent in Iodent than in Tropical germplasm, so the population difference reflects a combination of segregating variants rather than a single high- or low-expression allele.

The presence of three or more haplotypes each associated with different expression levels of the target gene is a potential mechanism to explain the observation that transcriptome-wide association studies (TWAS) frequently identify more and different genes associated with variation in a given phenotype than genome-wide association studies (GWAS) conducted using the same population and phenotype measurements (Torres-Rodríguez *et al*., 2024). For genes whose impact on phenotype acts via changes in transcript abundance, no single biallelic genetic marker can separate three distinct haplotypes, whereas direct measures of transcript abundance will reflect the impact of the complete set of different *cis*-regulatory variants present for the gene of interest in the study population.

Our study utilized expression data collected at a single time point from a single tissue. This had the virtue of limiting environmental and diurnal impacts on gene expression that can reduce power to identify genetic variants associated with gene expression, but it is unclear to what extent the specific variants we identify will translate to other tissues and other environments. One encouraging piece of evidence is that fine-mapped variants were more enriched in the set of accessible chromatin regions observed in both leaf and ear tissue than in open chromatin regions attributable to leaf alone. This is consistent with many of the *cis*-regulatory variants we observe impacting regulatory elements that act broadly rather than in a leaf-specific manner. In addition, it should be noted that several transcriptome-wide association studies have found that gene expression from one tissue or environment can identify true positive associations for phenotypes expressed in other tissues or scored in other environments (Li *et al*., 2021, 2024; Torres-Rodríguez *et al*., 2024).

In summary, resolving multiple signals offers a more informative representation of regulatory loci than reliance on single lead variants and identifies candidate alleles and haplotypes for functional investigation. While this study focused primarily on *cis*-regulatory variation, it remains to be seen whether the same patterns we observe are also present for *trans*-acting regulatory variation. This study provides a framework for interpreting expression diversity in crops by resolving the fine-scale architecture of *cis*-regulatory variation.

## Materials and Methods

### Plant material and sequencing data

The diversity panel consisted of 631 maize inbred lines from the Wisconsin Diversity Panel (Mazaheri *et al*., 2019). We used a previously published set of genome-wide variants generated from whole-genome resequencing data aligned to the B73 RefGen_v5 reference genome (Grzybowski *et al*., 2023; Hufford *et al*., 2021). Variant calls from the larger diversity panel were subset to the 631 lines included in this study and filtered using `bcftools` (Danecek *et al*., 2021). Variants with a minor allele frequency (MAF) below 1% were excluded, resulting in 29,322,258 variants available for subsequent analyses.

Transcriptomic data were obtained from a replicated field experiment conducted in 2020 at the University of Nebraska–Lincoln Havelock Farm near Lincoln, Nebraska, USA (40.852*^◦^* N, 96.616*^◦^* W). The experimental design, field management, and phenotypic measurements have been described previously (Mural *et al*., 2022; Sahay *et al*., 2023). Leaf tissue was collected 64 days after planting within a two-hour sampling interval and immediately flash frozen (Torres-Rodríguez *et al*., 2024). Transcript abundance was quantified using `kallisto` v0.46 (Bray *et al*., 2016) against the B73 RefGen_v5 primary-transcript reference (`primaryTranscriptOnly`), as described previously (Torres-Rodríguez *et al*., 2024).

All statistical analyses were performed in `R` (R Core Team, 2025). OpenAI Codex was used during code development to assist with code annotation, refactoring, and computational optimization.

### Population genetic analyses

Principal components of genome-wide genetic variation were calculated using `PLINK` v1.9 (Purcell *et al*., 2007) from variants with MAF *≥* 5% among the 631 lines.

Unbiased nucleotide diversity (*π*) was calculated for six subpopulations using `pixy` v1.2.7 in non-overlapping 100-bp windows (Korunes and Samuk, 2021), as described previously (Grzybowski *et al*., 2023). Input VCF files were generated using `GATK` v4.2.0.0 (Poplin *et al*., 2017) with the `-all-sites` option to retain records for both variable and invariant sites.

### Transcriptome diversity analysis

Gene-level abundance estimates from `kallisto` were imported into `R`, and lowly expressed genes were removed using `filterByExpr` from `edgeR` (Chen *et al*., 2025), leaving 18,197 genes. Expression values were normalized using the variance-stabilizing transformation (VST) implemented in `DESeq2` (Love *et al*., 2014) (Fig. S1). Additional details are provided in Supplementary Methods S1.

Because flowering time differs among maize genetic groups and may influence developmental state at sampling, its contribution to expression variation was removed before analyses of transcriptomic diversity (Mural *et al*., 2022). Flowering-time BLUPs were estimated across both experimental blocks using a mixed-effects model implemented in `lme4`, with inbred line fitted as a random effect (Bates *et al*., 2015). For each gene, VST-normalized expression was regressed on flowering time, and the residuals were used in subsequent analyses.

Principal component analysis (PCA) of flowering-time-adjusted expression was performed using `FactoMineR` (Lê *et al*., 2008). Concordance between genomic and transcriptomic population structure was assessed by canonical correlation analysis using the `cc` function from the `CCA` package (González and Déjean, 2023), jointly analyzing the first five genetic PCs and transcriptomic PCs derived from the expression PCA.

For each gene, the proportion of flowering-time-adjusted expression variation associated with genetic subpopulation was estimated using a linear model with subpopulation as a categorical predictor and summarized by *R*^2^. A null distribution was generated by randomly permuting subpopulation labels 1,000 times and recalculating gene-wise *R*^2^ values.

Within-subpopulation transcriptomic variability was quantified for each gene as the standard deviation of flowering-time-adjusted expression. Differences among subpopulations were tested using ANOVA based on linear mixed-effects model implemented in `lme4` (Bates *et al*., 2015), with subpopulation fitted as a fixed effect and gene identity as a random intercept.

### Identification of *cis* expression quantitative trait loci

*Cis*-expression quantitative trait locus (eQTL) mapping was performed using log_2_-transformed counts per million (log_2_-CPM) gene-expression values calculated with `edgeR` (Chen *et al*., 2025) as quantitative phenotypes. Association mapping was conducted using a mixed linear model (MLM) implemented in OmicS-data-based Complex Trait Analysis (`OSCA`; Zhang *et al*., 2019). For each gene, variants located within *±*2 Mb of the transcription start site (TSS) were tested for association with gene expression.

To account for latent sources of expression variation, PCA was performed on the gene-expression matrix, and the first 10 expression PCs were included as covariates (Supplementary Methods S2; Fig. S2). Potential redundancy between expression PCs and known biological and genetic covariates was evaluated by regressing flowering time and the first five genetic PCs against the expression PCs. Genetic PCs strongly explained by expression PCs (*R*^2^ *>* 0.9) were excluded to avoid redundant adjustment. Flowering time and the remaining genetic PCs were retained together with the 10 expression PCs, resulting in 15 fixed-effect covariates. A genomic relationship matrix calculated using `GCTA` (Yang *et al*., 2011) was included as a random effect to account for genetic relatedness among lines. Details of covariate selection and evaluation are provided in Supplementary Methods S2.

Multiple testing was controlled using Independent Hypothesis Weighting (IHW) (Ignatiadis *et al*., 2016), with the absolute distance between each tested variant and the TSS of its target gene used as the informative covariate (Fig. S3). Significant *cis*-eQTL associations were identified at a false discovery rate (FDR) of 1%. Details of the IHW implementation, including construction of the TSS-distance covariate, are provided in Supplementary Methods S3.

To evaluate the effect of sample size on eQTL discovery, we performed a downsampling analysis using the same set of 18,197 expressed genes. Eight subsets of the full diversity panel were generated by stratified random sampling while approximately preserving the relative representation of the major genetic subpopulations. Within each subset, variants with MAF *<* 5% were excluded, resulting in approximately 15.5 million variants available for association testing at each sample size.

For the downsampling analysis, association testing was performed using the general linear model (GLM) implemented in `OSCA`. The GLM was used because mixed-model analyses showed an increasing frequency of convergence failures at smaller sample sizes. The first five genetic PCs, calculated separately within each downsampled dataset, were included as covariates. To provide a consistent significance criterion across sample sizes, associations were considered significant at a Bonferroni-corrected family-wise error rate of 5%. The number of genes with at least one significant *cis*-eQTL was quantified at each sample size to characterize the relationship between sample size and eQTL discovery.

### Fine-mapping and estimation of allelic effect sizes

Fine-mapping of *cis*-eQTL signals was performed using the `susieR` package (Wang *et al*., 2020b). For each eGene, all variants within the same *±*2 Mb window around the TSS used for the initial eQTL mapping were included. SuSiE models the association signal as a sum of single effects, allowing multiple distinct regulatory signals to be identified within the same locus. For each inferred signal, a 95% credible set was constructed containing variants that collectively accounted for at least 95% posterior probability of containing the causal variant for that effect.

Before fine-mapping, normalized log_2_-CPM expression values and genotype dosages were adjusted for the same fixed-effect covariates used in the eQTL analysis. Covariate effects were removed from both the expression vector and genotype matrix, and the resulting residuals were used as input to `susieR`. Models were fitted with a maximum of 10 single-effect components per gene, residual variance estimated from the data, and a maximum of 500 iterations. Credible sets were constructed at 95% coverage, and sets failing the default SuSiE purity criterion (minimum absolute within-set correlation of 0.5) were excluded. Posterior inclusion probabilities (PIPs) were estimated for individual variants. For downstream analyses requiring a single representative variant per signal, the variant with the highest PIP within each credible set was selected as the lead fine-mapped eQTL variant.

The magnitude of regulatory effects was quantified using allelic fold change (aFC), which represents the multiplicative change in gene expression associated with the alternative allele relative to the reference allele. Because many eGenes contained multiple independently fine-mapped *cis*-regulatory signals, effect sizes were estimated using the aFC-n framework (Ehsan *et al*., 2024), which extends aFC estimation to jointly model multiple regulatory variants affecting the same gene. aFC-n was applied using the implementation available at https://github.com/PejLab/aFCn, with gene expression adjusted for the same covariates used in the fine-mapping analysis. Effect sizes were expressed as log_2_(aFC), with positive and negative values indicating increased and decreased expression associated with the alternative allele, respectively.

### Expression heritability and multi-signal prediction

Narrow-sense heritability of gene expression was estimated for individual eGenes using a genomic relationship matrix and a linear mixed model implemented in the `sommer` package (Giovanny, 2016). For each gene, expression was modeled with a random additive genetic effect whose covariance structure was defined by the genomic relationship matrix and an independent residual component. Narrow-sense heritability was calculated as

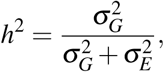

where *σ_G_* ^2^ and *σ_E_* ^2^ denote the additive genetic and residual variance components, respectively. Heritability was estimated from both unadjusted log_2_-CPM expression and expression adjusted for the same fixed-effect covariates used in the eQTL analysis. Covariate-adjusted estimates were used for downstream comparisons with the number of fine-mapped regulatory signals.

To determine whether multiple independently fine-mapped *cis*-regulatory signals improved prediction of gene expression relative to a conventional single-variant model, the 631 lines were divided into training and test sets. A common split was used for all genes, with 80% of lines assigned to the training set and 20% to the test set. Sampling was stratified by genetic subpopulation to preserve their relative representation in both sets.

For each eGene, two genetic prediction models were compared. The single-variant model contained the variant with the lowest marginal *p*-value from the `OSCA` *cis*-eQTL analysis. The multi-signal model contained one representative fine-mapped variant from each credible set, defined as the variant with the highest PIP within that set. When multiple credible sets were detected, their representative variants were fitted jointly in a multiple linear regression model. Model coefficients were estimated exclusively in the training set and subsequently used to predict covariate-adjusted expression in the held-out test set.

Predictive performance was quantified as

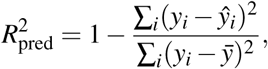

where *y_i_* and *y*^*_i_* are the observed and predicted covariate-adjusted expression values, respectively, in the test set. Test-set *R*^2^ was compared between the single-variant and multi-signal models to evaluate whether multiple independently fine-mapped *cis*-regulatory signals improved prediction of gene expression.

### Functional annotation and regulatory enrichment

Fine-mapped *cis*-eQTL signals, represented by the variant with the highest PIP within each credible set, were functionally annotated relative to the B73 RefGen_v5 gene annotation. Genomic overlaps were determined using the `GenomicRanges` and `GenomicFeatures` Bioconductor packages (Lawrence *et al*., 2013). Variants were classified according to overlap with 5*^′^* UTRs, 3*^′^* UTRs, coding sequences (CDS), promoters, introns, or intergenic regions. Promoters were defined as regions extending 2 kb upstream and 200 bp downstream of transcription start sites.

Positional enrichment of lead fine-mapped *cis*-eQTL variants was evaluated relative to significant associations identified in the primary *cis*-eQTL analysis. Absolute distances between variants and the transcription start sites (TSSs) of their associated genes were grouped into the same distance bins for the fine-mapped lead variants and the significant association background.

Enrichment of fine-mapped variants within genomic annotations was evaluated using a matched permutation framework. For each fine-mapped signal, control variants were sampled from the same *±*2 Mb *cis* region surrounding the TSS of the corresponding target gene and matched for MAF, absolute distance from the target-gene TSS, and variant type (SNP or indel). Fold enrichment was calculated relative to the expected overlap obtained from 5,000 matched permutations, and empirical *p*-values were corrected for multiple testing using the Benjamini–Hochberg procedure.

Regulatory enrichment was evaluated using published maize chromatin-accessibility and transcription-factor-binding datasets (Ricci *et al*., 2019; Tu *et al*., 2020). BED files containing leaf ATAC-seq peaks and transcription-factor ChIP-seq peaks were obtained from MaizeGDB (Woodhouse *et al*., 2025). Fine-mapped variants were intersected with leaf ATAC-seq peaks and replicate-supported ChIP-seq consensus peaks for 104 maize transcription factors. Enrichment within accessible chromatin was assessed using controls matched for MAF, TSS distance, and variant type. For transcription-factor-binding analyses, controls were additionally matched for overlap with leaf ATAC-seq peaks, allowing enrichment for individual transcription factors to be evaluated conditional on local chromatin accessibility. Full annotation definitions, matching procedures, and permutation tests are described in Supplementary Methods S4.

### Population differentiation and evolutionary constraint of regulatory variation

To investigate the relationship between population differentiation and gene-expression divergence, differential expression and variant-level *F*_ST_ were evaluated across matched population contrasts. Differential expression was analyzed for the same set of expressed genes used for *cis*-eQTL mapping using `limma-voom` (Ritchie *et al*., 2015). Genes with Benjamini–Hochberg-adjusted *p*-values *<* 0.05 were considered differentially expressed.

Population contrasts included Stiff Stalk (SS), Non-Stiff Stalk (NSS), Iodent, and Tropical germplasm, as well as comparisons between older lines (era I; *≤* 2000) and more recently released lines (era II; *>* 2000). Differentiation between SS and NSS was evaluated separately within eras I and II to assess whether divergence between the two major heterotic groups differed across breeding history. Per-variant *F*_ST_ values were calculated for the corresponding population contrasts using `GCTA` (Yang *et al*., 2011). Fine-mapped eQTL variants were subsequently annotated with both the adjusted *p*-value from the differential-expression analysis of their associated eGene and the variant-level *F*_ST_ for the corresponding population contrast. The relationship between transcriptional divergence and regulatory differentiation was evaluated by grouping differentially expressed genes into deciles according to *−* log_10_-transformed adjusted *p*-values and comparing the *F*_ST_ distributions of their fine-mapped eQTL variants across deciles.

To evaluate the relationship between evolutionary constraint and *cis*-regulatory variation, Genomic Evolutionary Rate Profiling (GERP) scores were obtained from the B73 RefGen_v5 comparative-genomics resource (Davydov *et al*., 2010; Hufford *et al*., 2021). Only sites with positive GERP scores were retained. For each annotated gene, the mean positive GERP score across the gene body was calculated. Genes in the upper 10% of the genome-wide distribution were classified as highly constrained, with all remaining genes used as the comparison group.

The magnitude and complexity of *cis*-regulatory variation were compared between highly constrained and other genes. Absolute fine-mapped effect sizes, expressed as *|* log_2_(aFC)*|*, were compared using a Wilcoxon rank-sum test. Differences in the number of SuSiE credible sets per eGene were evaluated using a quasi-Poisson generalized linear model with a log link to account for overdispersion in credible-set counts. These analyses tested whether genes under stronger evolutionary constraint exhibit smaller regulatory effect sizes or fewer independently fine-mapped *cis*-regulatory signals.

## Acknowledgments

This work was supported by Grant 2023/49/B/NZ9/00766 from the National Science Centre (NCN), Poland.

## Author contributions

MWG and JCS conceived of the experiments and analyses. MWG conducted the experiments and analyses, interpreted the results and drafted the manuscript. JCS and MWG both revised and approved the final version of the manuscript.

## Data availability

All of the scripts used in this study are available at: https://github.com/mgrzy/Maize_Transcriptome_ `Diversity_v2`. Code and processed gene expression matrices and QTL mapping results are available from Zenodo (https://doi.org/10.5281/zenodo.18670486).

## Supplementary Data

**Supplementary Methods S1–S4.** Expression normalization (S1), selection of covariates for *cis*-eQTL mapping (S2), implementation of Independent Hypothesis Weighting (S3), and functional annotation definitions, matching procedures, and permutation tests for enrichment analyses (S4).

**Table S1.** Results of multivariate analyses integrating genotype, transcriptomic, and metadata variables (XLSX).

**Table S2.** Fine-mapping results for lead *cis*-eQTNs, including credible sets and posterior inclusion probabilities (XLSX).

**Table S3.** Allelic fold change (aFC) estimates for lead *cis*-eQTNs (XLSX).

**Figure S4.** Genetic principal component analysis of WiDiv lines in the context of broader *Zea* diversity.

**Figure S5.** Summary of *cis*-eQTL discovery: number of associated SNPs per eGene, distance from the target-gene transcription start site, and minor allele frequency distribution.

**Figure S6.** Effect of sample size on eGene discovery and the mean number of significant SNPs per gene, assessed by stratified downsampling of the WiDiv panel.

**Figure S7.** Comparison of narrow-sense gene-expression heritability estimated from unadjusted and covariate-adjusted expression.

**Figure S8.** Relationships of expression mean and standard deviation with narrow-sense expression heritability and the number of fine-mapped regulatory signals.

**Figure S9.** Comparison of evaluation-set performance between single-variant and fine-mapped multi-variant expression models.

**Figure S10.** Enrichment of fine-mapped *cis*-eQTL variants in consensus ChIP-seq peaks for 104 maize transcription factors, using standard matched controls and controls additionally matched for leaf chromatin accessibility.

**Figure S1.** Effect of expression normalization on the relationship between gene-wise mean expression and standard deviation: raw counts, TMM-normalized log_2_-CPM, and variance-stabilized expression.

**Figure S2.** Selection of expression principal components as covariates for *cis*-eQTL mapping.

**Figure S3.** Evaluation of transcription-start-site distance and minor allele frequency as covariates for Independent Hypothesis Weighting in the *cis*-eQTL analysis.

