## Supplementary Methods and Figures for "Haplotype-rich *cis*-regulation underlies transcriptomic diversity across the breeding history of maize (*Zea mays*)"

**References** **6**

**Supplementary figures** **7**

### Supplementary methods

#### Supplementary Methods S1: Expression normalization and variance stabilization

Gene-level RNA-seq counts were restricted to samples that could be matched to genotyped Wisconsin Diversity Panel lines. Lowly expressed genes were removed using `edgeR::filterByExpr`. Library-size normalization factors were subsequently estimated using the trimmed mean of M-values (TMM) method, and expression values were converted to $\log_2$ counts per million ($\log_2$-CPM) using `edgeR`[1], with a prior count of 2[1]. The $\log_2$ transformation compresses the dynamic range of RNA-seq expression values, reducing the disproportionate influence of highly expressed genes and partially stabilizing the relationship between the expression mean and variance (Fig. S1 A and B). The resulting filtered $\log_2$-CPM matrix was used as the expression phenotype for the primary *cis*-eQTL analysis.

For analyses of transcriptome-wide expression diversity, counts from the same filtered gene set were transformed using the variance-stabilizing transformation (VST) implemented in `DESeq2`[2]. In contrast to a simple logarithmic transformation, VST uses the estimated mean–dispersion relationship of the RNA-seq count data to transform expression values such that their variance is approximately independent of their mean. This is particularly useful for comparisons of expression variability across genes and samples because it reduces the strong dependence of variance on expression level that is characteristic of count data (Fig. S1 C).

The VST-transformed expression matrix was therefore used for transcriptome-diversity analyses, including comparisons of genome-wide expression variation among genetic groups.

As a normalization diagnostic, the relationship between gene-wise mean expression and standard deviation was compared among raw counts, TMM-normalized $\log_2$-CPM values, and VST-transformed values. Both $\log_2$-CPM and VST substantially reduced the strong mean–variance dependence observed for raw counts, with VST providing the expression scale used for analyses explicitly focused on transcriptomic variability (Fig. S1).

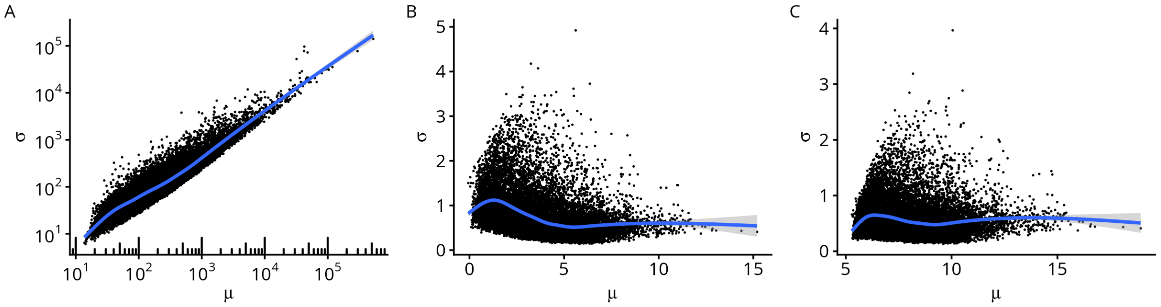

Supplementary Figure S1: Effect of expression normalization on the gene-wise mean–variance relationship. Gene-wise mean expression and standard deviation are shown for (A) raw counts, (B) TMM-normalized log2-CPM values used for the primary eQTL analysis, and (C) Variance-stabilized expression values. Smooth curves summarize the mean–variance relationship.

### Supplementary Methods S2: Selection of covariates for *cis*-eQTL mapping

Latent expression covariates were selected by principal component analysis (PCA) of the filtered $\log_2$-CPM expression matrix. Expression values were centered and scaled prior to PCA. The number of expression principal components (PCs) was initially evaluated using the elbow and Buja–Eyuboglu procedures implemented in `PCAForQTL`[3].

The elbow method suggested inclusion of 25 expression PCs, whereas the Buja–Eyuboglu procedure identified 68 PCs (Fig. S2). However, these procedures select latent covariates independently of the genetic relationship matrix (GRM) subsequently incorporated as a random effect in the mixed linear model (MLM). When 25 expression PCs were included together with the GRM, approximately 20% of genes produced model-convergence failures. We therefore evaluated a reduced number of expression covariates and found that inclusion of 10 expression PCs allowed successful MLM fitting for more than 99% of genes while still accounting for major latent sources of transcriptomic variation. Consequently, the first 10 expression PCs were included as covariates in the primary *cis*-eQTL analysis.

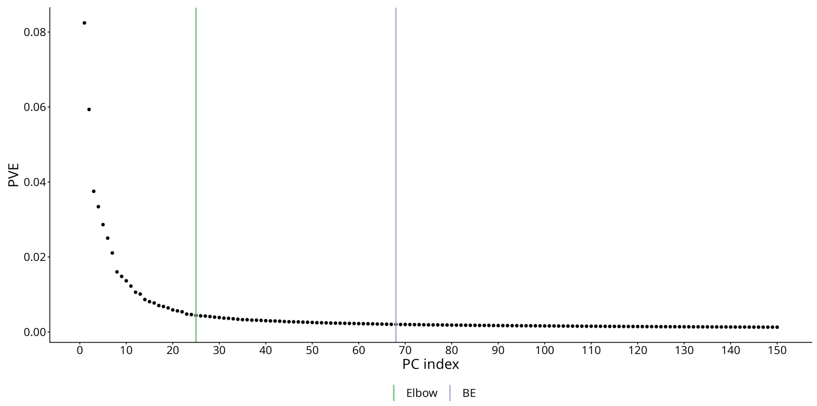

Supplementary Figure S2: Selection of expression principal components for cis-eQTL mapping. The scree plot summarizes variance captured by expression PCs and the PC-number estimates obtained using the elbow and Buja–Eyuboglu procedures. Ten expression PCs were retained in the primary eQTL model.

### Supplementary Methods S3: IHW diagnostics and multiple-testing adjustment

Absolute distance between each tested variant and the transcription start site (TSS) of its target gene was evaluated as an informative covariate for Independent Hypothesis Weighting (IHW)[4]. Minor allele frequency (MAF) was also considered as a candidate covariate. The relationship between each candidate covariate and association strength was first examined across all tested variant–gene pairs (Fig. S3A,B). Strong associations were markedly enriched close to the TSS, with the excess of small *P*-values progressively decreasing with increasing TSS distance (Fig. S3A). In contrast, MAF showed a less pronounced monotonic relationship with association strength (Fig. S3B).

For further evaluation of TSS distance, the 2-Mb *cis* interval was divided into ten 200-kb bins. The number of tested variant–gene pairs was similar across distance bins, ranging from approximately 90.6 to 93.0 million tests per bin (Fig. S3C). The corresponding *P*-value distributions were then examined separately within each distance stratum (Fig. S3D). Proximal bins showed a substantially greater excess of very small *P*-values, consistent with a higher frequency of true regulatory associations near the TSS, whereas the non-significant portion of the *P*-value distribution was broadly comparable across distance strata. Together, these diagnostics supported absolute TSS distance as an informative and appropriately ordered covariate for IHW, and it was therefore used in the primary analysis.

Because the complete *cis*-eQTL scan comprised approximately $9.2 \times 10^8$ association tests, IHW was applied using a truncated *P*-value representation to reduce memory requirements. Associations with $P \leq 10^{-3}$ were retained explicitly, while the total number of hypotheses in each of the ten TSS-distance bins was calculated from the complete OSCA[5] output and supplied to IHW. This approach retained the correct number of hypotheses represented by each covariate stratum without requiring storage of the large number of clearly non-significant association results.

IHW was performed at a target false discovery rate (FDR) of 1%, with TSS-distance bins treated as an ordinal covariate. Data-driven hypothesis weights were estimated using fivefold cross-validation with a fixed random seed of 1 and subsequently incorporated into the weighted Benjamini–Hochberg procedure. Variant–gene associations with IHW-adjusted $P < 0.01$ were considered significant *cis*-eQTLs.

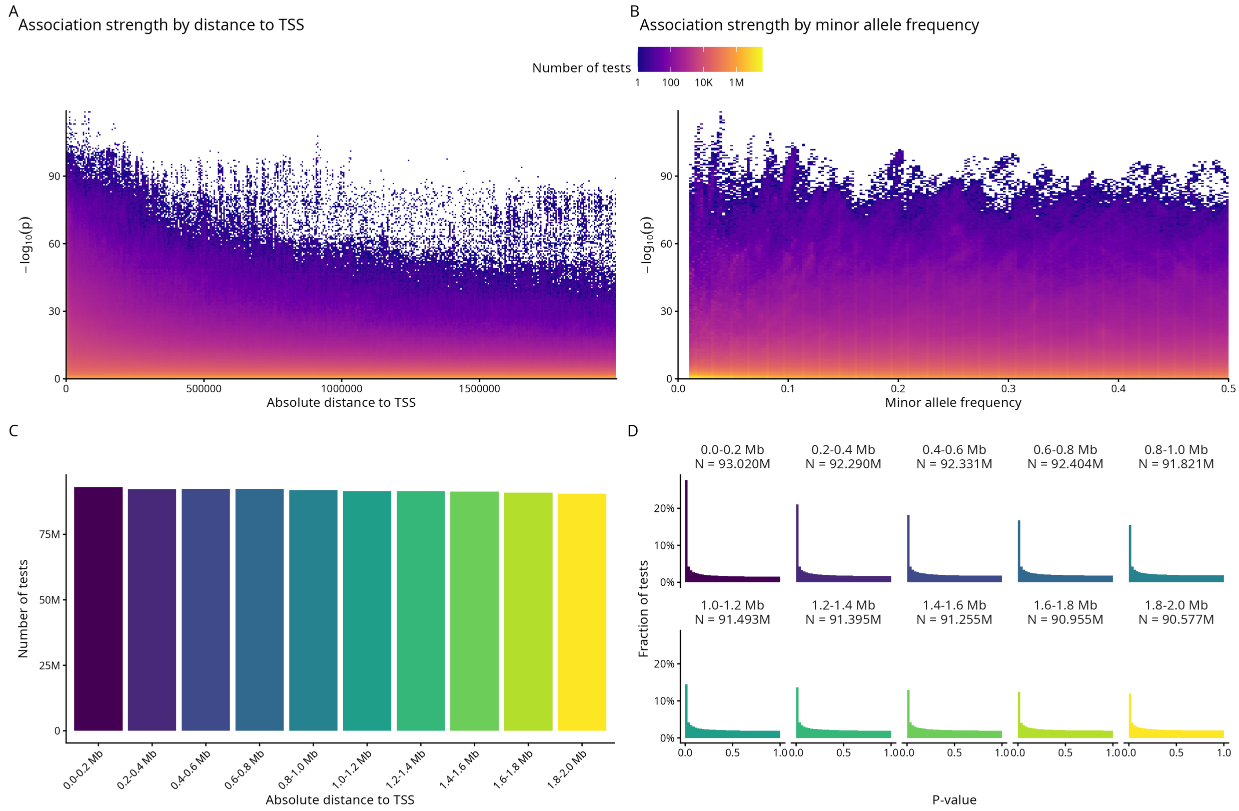

Supplementary Figure S3: Selection of the covariate for Independent Hypothesis Weighting in the cis-eQTL analysis. (A) Relationship between absolute variant distance to the transcription start site (TSS) and association significance. (B) Relationship between minor allele frequency (MAF) and association significance. Color indicates the density of association tests. (C) Number of variant–gene association tests within ten 200-kb TSS-distance bins across the 2-Mb cis interval. (D) P-value distributions within the same TSS-distance bins. Proximal variants show a greater excess of small P-values, whereas the non-significant portion of the distribution remains broadly similar among bins, supporting TSS distance as an informative covariate for IHW.

### Supplementary Methods S4: Functional annotation and regulatory enrichment

#### Genomic annotation of fine-mapped eQTL variants

Functional annotation was performed using one lead variant from each credible set, defined as the variant with the highest posterior inclusion probability (PIP) within that set. Gene models were obtained from the B73 RefGen_v5 annotation (Zm-B73-REFERENCE-NAM-5.0_Zm00001eb.1) and imported into R using the `GenomicFeatures`, `txdbmaker`, and `GenomicRanges` Bioconductor packages[6].

Gene bodies, transcripts, exons, coding sequences (CDS), 5′ untranslated regions (5′ UTRs), 3′ untranslated regions (3′ UTRs), and introns were derived directly from the genome annotation. Promoter regions were defined relative to transcript-specific TSSs as extending 2,000 bp upstream and 200 bp downstream of the TSS. Overlaps between fine-mapped variants and genomic features were calculated without regard to strand.

Because individual genomic positions can overlap multiple annotations, all binary feature-overlap indicators were retained. In addition, each variant was assigned a single primary genomic annotation for enrichment analyses using the following hierarchical priority:

$$5'\mathrm{UTR} > 3'\mathrm{UTR} > \mathrm{CDS} > \text{other exon} > \text{promoter} > \text{intron} > \text{intergenic}.$$

Thus, for example, a variant overlapping both a promoter and a 5′ UTR was assigned to the 5′ UTR category for analyses requiring mutually exclusive annotations.

#### Matched genomic-background enrichment

Enrichment of fine-mapped eQTL variants within genomic annotation classes was evaluated relative to matched control variants drawn from the complete set of genotyped variants. Controls were restricted to the same ±2 Mb *cis* window around the TSS of the corresponding target eGene used for eQTL mapping and fine-mapping.

For each fine-mapped signal, candidate control variants were matched according to target gene, variant type (SNP or indel), MAF, and absolute distance from the target-gene TSS. MAF was divided into intervals of width 0.05. Absolute TSS distance was divided into the following intervals: 0–1 kb, 1–2 kb, 2–5 kb, 5–10 kb, 10–25 kb, 25–50 kb, 50–100 kb, 100–250 kb, 250–500 kb, 0.5–1 Mb, and 1–2 Mb. Upstream and downstream variants were treated equivalently on the basis of absolute TSS distance.

The fine-mapped lead variant itself was excluded from its control pool. To limit computational requirements while retaining a large candidate set, at most 500 candidate variants were retained per matching stratum. Fine-mapped signals with fewer than five eligible matched controls were excluded from the genomic-context permutation analysis.

A total of 5,000 permutations was performed. During each permutation, one control variant was sampled independently from the matched pool of every eligible fine-mapped signal. The number of sampled variants belonging to each genomic annotation class was recorded. The expected number of variants in each category was calculated as the mean count across permutations, and fold enrichment was calculated as

$$\text{Fold enrichment} = \frac{N_{\text{observed}}}{\text{mean}(N_{\text{permuted}})}.$$

One-sided empirical probabilities for enrichment and depletion were calculated as

$$P_{\text{enrichment}} = \frac{1 + \sum(N_{\text{permuted}} \geq N_{\text{observed}})}{1 + N_{\text{permutations}}},$$

and

$$P_{\text{depletion}} = \frac{1 + \sum(N_{\text{permuted}} \leq N_{\text{observed}})}{1 + N_{\text{permutations}}}.$$

Two-sided empirical *p*-values were calculated as twice the smaller of the enrichment and depletion probabilities, with values constrained to a maximum of 1. Multiple testing across genomic annotation categories was controlled using the Benjamini–Hochberg false discovery rate procedure.

#### Chromatin-accessibility enrichment

BED files containing published maize chromatin-accessibility peaks were downloaded from the MaizeGDB website[7,8]. The downloaded peak coordinates corresponded to the B73 RefGen_v5 genome assembly. Chromosome identifiers were standardized before analysis, and BED coordinates were converted from the 0-based, half-open BED convention to the 1-based coordinate system used for variant positions. Fine-mapped variant positions were intersected with leaf ATAC-seq peaks using `GenomicRanges`.

Enrichment within accessible chromatin was assessed using the same general matched-permutation strategy as for genomic annotations. Candidate control variants were drawn from the ±2 Mb *cis* region of the corresponding target gene and matched according to MAF, absolute TSS distance, and variant type. The same MAF and TSS-distance bins described above were used, and fine-mapped variants were excluded from the control pool.

For each of 5,000 permutations, one eligible matched control was sampled for every fine-mapped variant, and overlap with ATAC-seq peaks was recorded. Fold enrichment and empirical $p$-values were calculated from the resulting permutation distribution.

#### Transcription-factor ChIP-seq enrichment

BED files containing published maize transcription-factor ChIP-seq peaks were downloaded from the MaizeGDB website[8,9]. The dataset comprised replicate-supported consensus peaks for 104 maize transcription factors represented in B73 RefGen_v5 coordinates. Chromosome identifiers and BED coordinates were standardized as described above, and enrichment was evaluated independently for each transcription factor.

Because fine-mapped eQTL variants were themselves enriched within accessible chromatin, transcription-factor-binding enrichment was tested conditional on general chromatin accessibility. Candidate control variants were therefore matched to each fine-mapped variant according to MAF, absolute distance from the target-gene TSS, variant type, and binary overlap status with leaf ATAC-seq peaks. Consequently, an ATAC-positive fine-mapped variant was compared only with ATAC-positive controls, whereas an ATAC-negative variant was compared with ATAC-negative controls. Exact matching used MAF bins of width 0.05 and the TSS-distance intervals described above.

For each transcription factor, 5,000 permutations were performed by independently sampling one matched control for every eligible fine-mapped variant. The number of variants overlapping ChIP-seq peaks for the corresponding transcription factor was calculated for each permutation and compared with the observed overlap. Fold enrichment and empirical $p$-values were calculated from these permutation distributions. Multiple testing across transcription factors was controlled using the Benjamini–Hochberg false discovery rate procedure.

### Supplementary figures

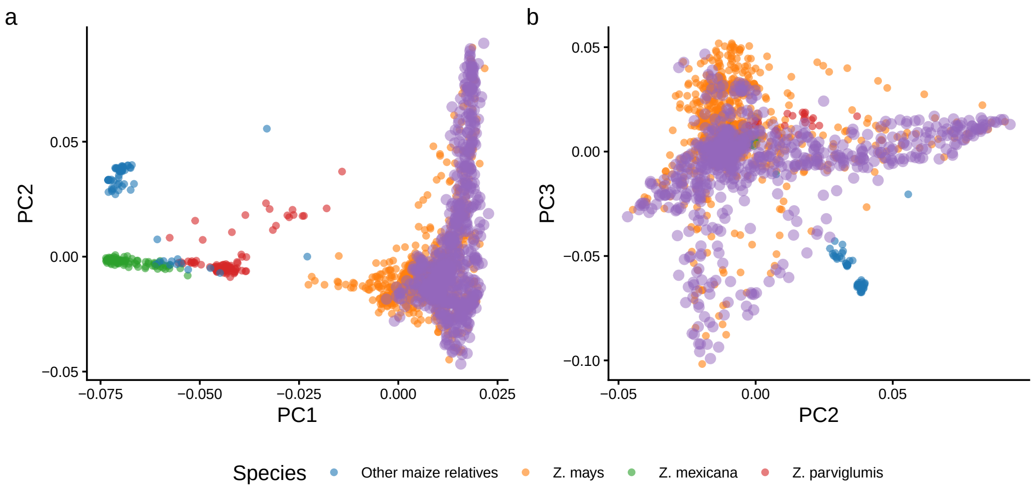

Supplementary Figure S4: Genetic principal component analysis of WiDiv lines in the context of broader Zea diversity. (A) PC1 versus PC2 and (B) PC2 versus PC3. WiDiv accessions are highlighted relative to maize, teosinte, and other maize relatives in the broader diversity panel.

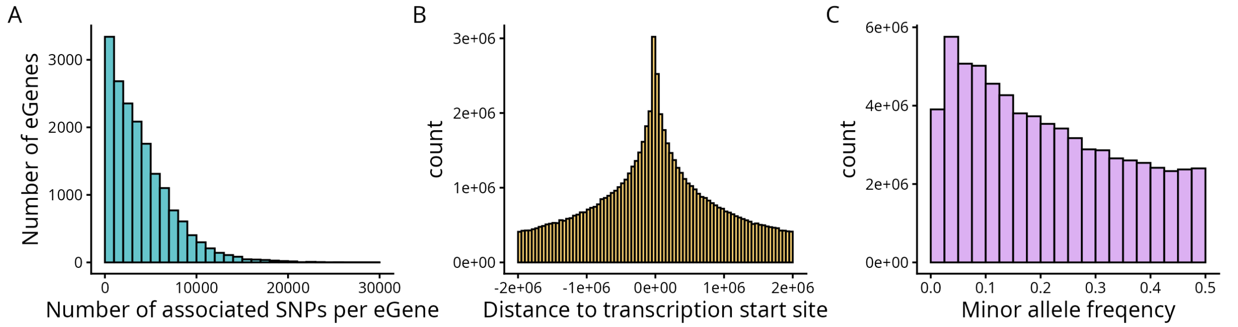

Supplementary Figure S5: Summary of cis-eQTL discovery in the WiDiv panel. (A) Distribution of the number of associated SNPs per eGene. (B) Distribution of distances between associated SNPs and the transcription start site of their target gene. (C) Minor allele frequency distribution of associated SNPs.

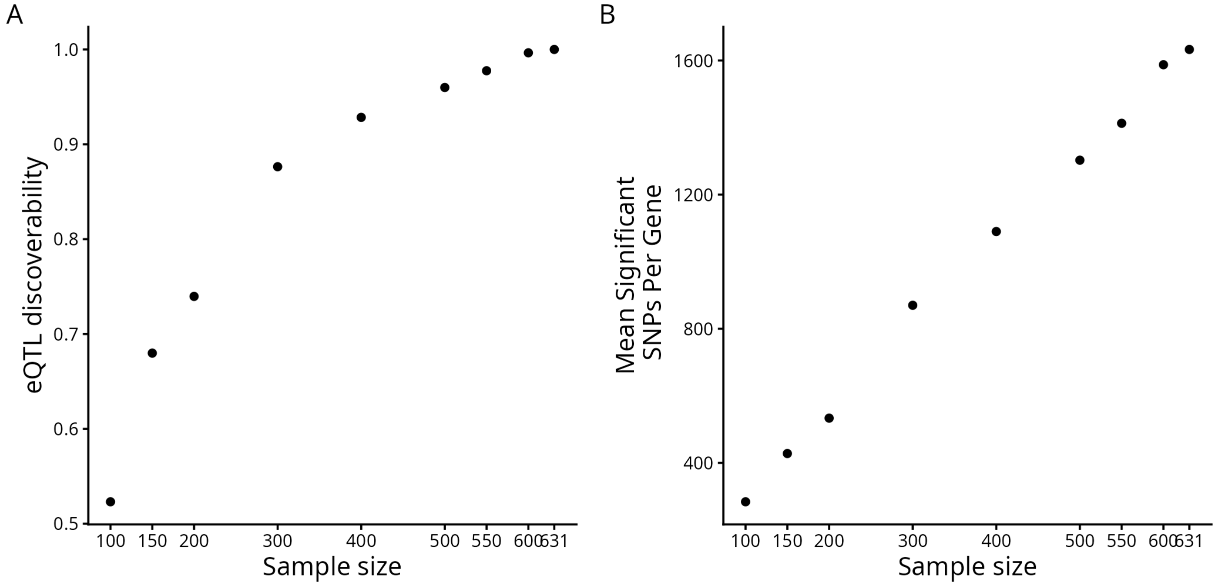

Supplementary Figure S6: Effect of sample size on cis-eQTL discovery. Stratified downsampling of the WiDiv panel was used to quantify the number of genes with at least one significant cis-eQTL across progressively larger sample sizes.

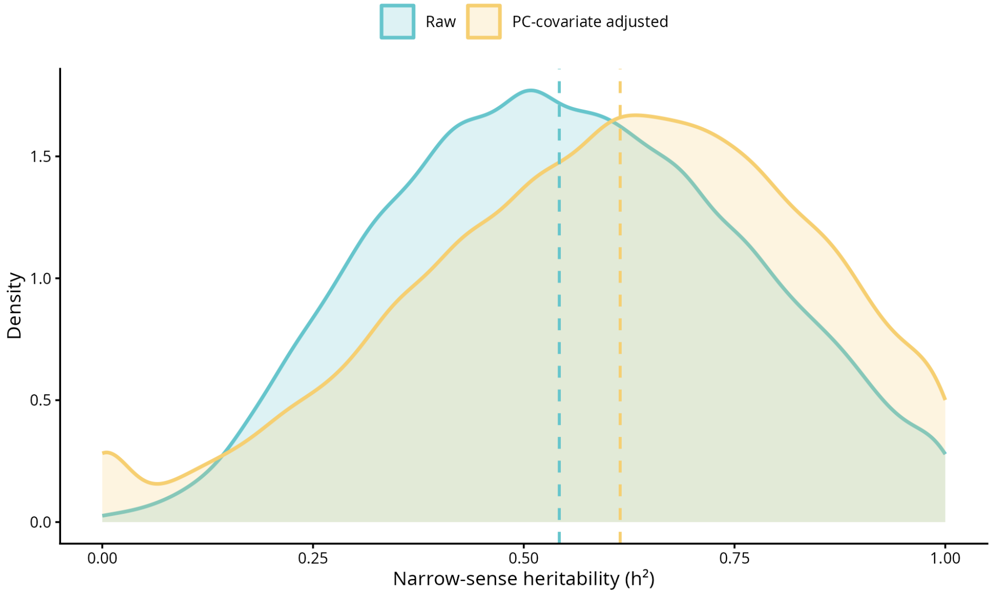

Supplementary Figure S7: Comparison of narrow-sense heritability estimates of gene expression calculated from raw and covariate-adjusted expression values.

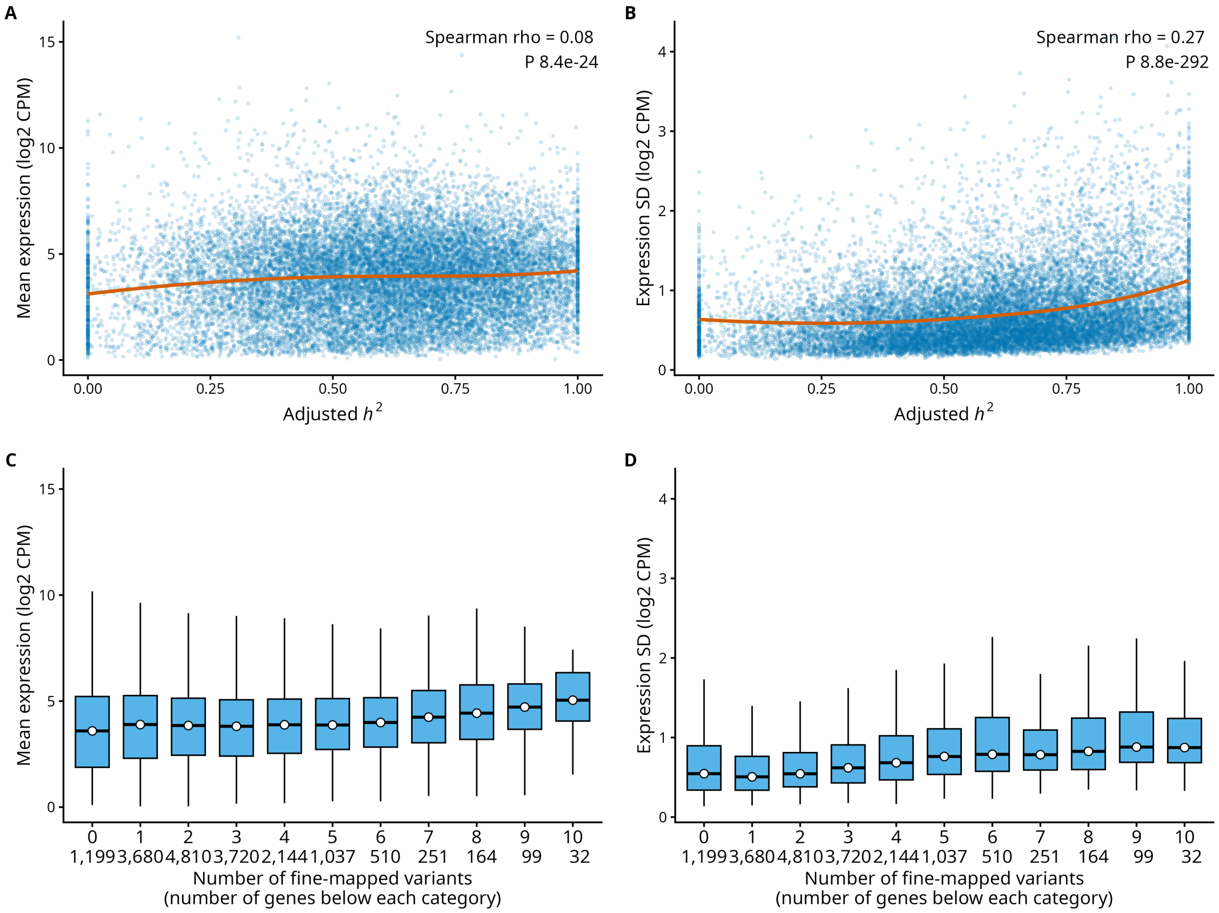

Supplementary Figure S8: Relationship between expression characteristics, narrow-sense expression heritability, and the number of fine-mapped regulatory signals. (A,B) Mean and standard deviation of log2-CPM expression as a function of covariate-adjusted narrow-sense heritability. (C,D) Mean and standard deviation of log2-CPM expression according to the number of fine-mapped SuSiE lead variants per gene.

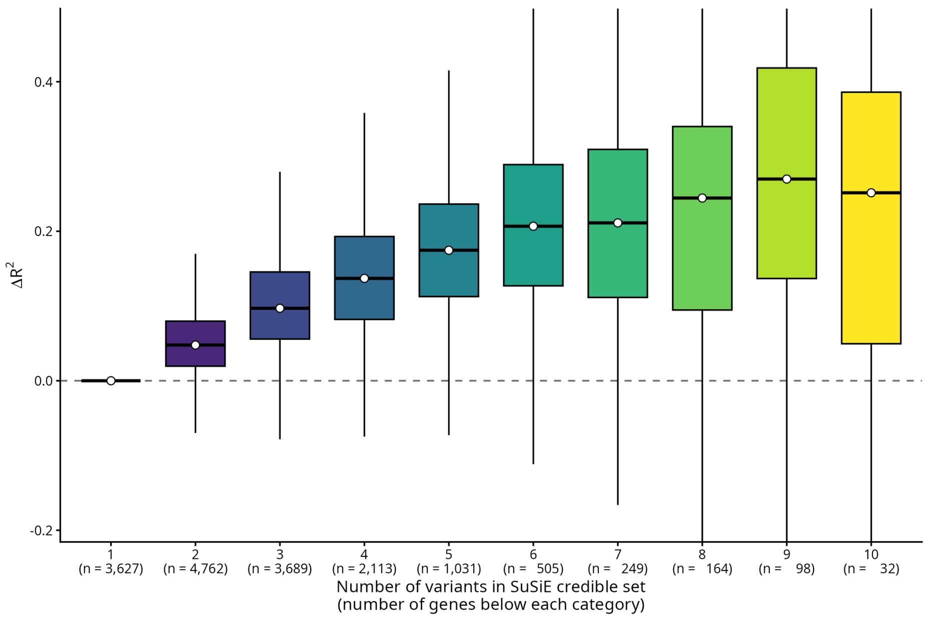

Supplementary Figure S9: Difference in held-out prediction performance between fine-mapped multi variants models and single-variant models across SuSiE credible-set sizes. Delta R2 is calculated as multi variants model test R2 minus single-variant models test R2; positive values indicate better held-out prediction by the fine-mapped multi variants model model.

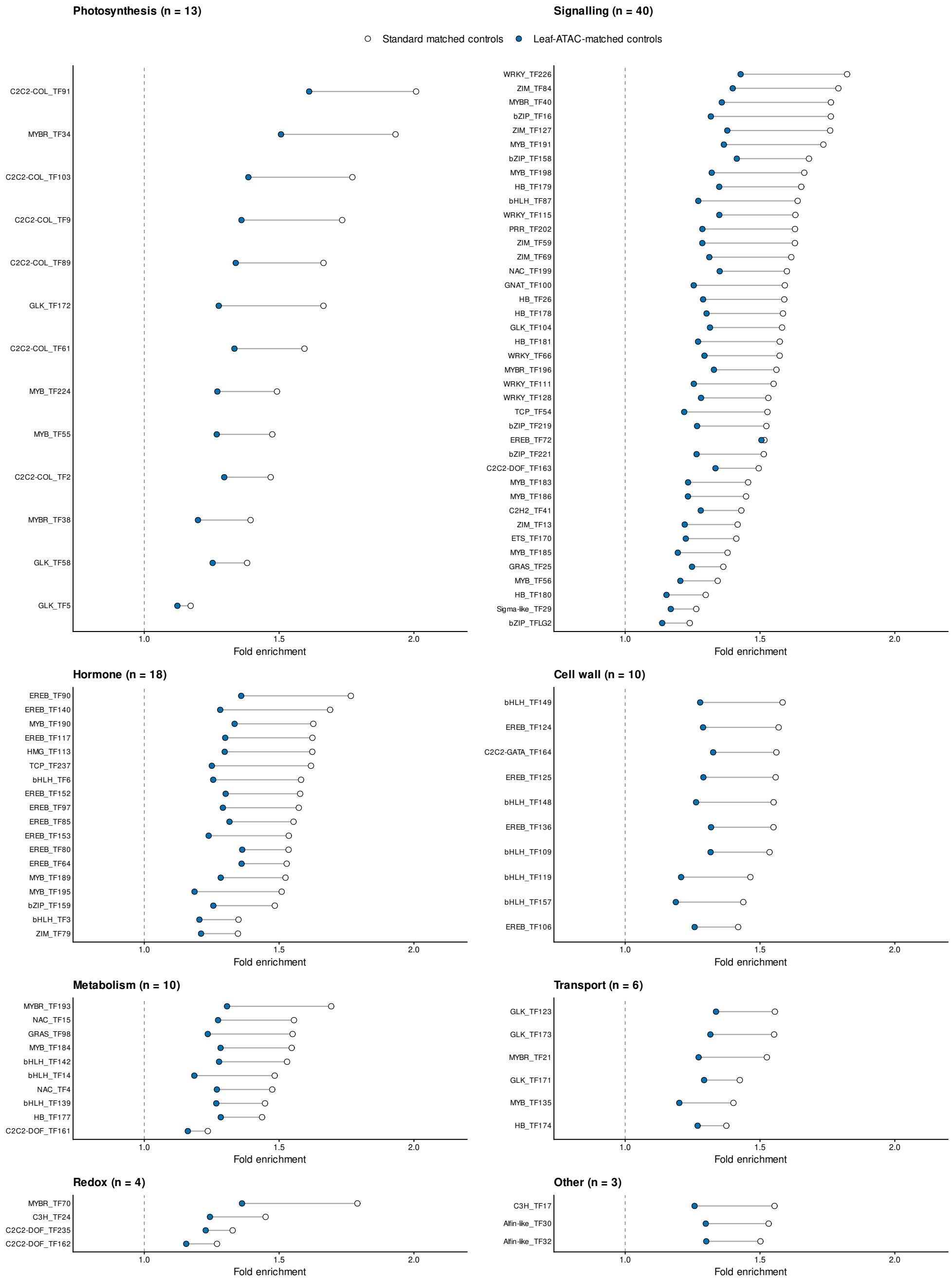

Supplementary Figure S10: Fine-mapped cis-eQTL enrichment in consensus ChIP-seq peaks for all 104 maize transcription factors. Transcription factors are grouped according to the eight functional paper classes used in Figure 3. Open points show enrichment relative to standard matched controls; blue points show enrichment relative to controls additionally matched for leaf chromatin accessibility. Lines connect estimates for the same transcription factor; the dashed line denotes no enrichment.
